# Lpnet: Reconstructing Phylogenetic Networks from Distances Using Integer Linear Programming

**DOI:** 10.1101/2021.10.08.463657

**Authors:** Mengzhen Guo, Stefan Grünewald

**Affiliations:** Shanghai Institute of Nutrition and Health, University of Chinese Academy of Sciences, Chinese Academy of Sciences, Shanghai 200031, PR China

**Keywords:** Phylogenetic networks, Neighbor-net, Integer linear programming, Circular split systems, Distance-based phylogenetics

## Abstract

We present Lpnet, a variant of the widely used Neighbor-net method that approximates pairwise distances between taxa by a circular phylogenetic network. We first apply standard methods to construct a binary phylogenetic tree and then use integer linear programming to compute an optimal circular orderings that agrees with all tree splits. This approach achieves an improved approximation of the input distance for the clear majority of experiments that we have run for simulated and real data. We release an implementation in R that can handle up to 94 taxa and usually needs about one minute on a standard computer for 80 taxa. For larger taxa sets, we include a top-down heuristic which also tends to perform better than Neighbor-net.

## Introduction

Neighbor-net is a widely used distance-based method that approximates the input distance by (the distance induced by) a weighted circular split system on the same taxa set, which is then visualized as a splits graph. It has been used to analyze numerous real data sets, and it is a crucial step of other phylogenetic tools, for example to construct coalescent-based phylogenetic networks (Allman *et al*., 2019), to detect and filter out homoplastic sites of a sequence alignment (Dress *et al*., 2008), and to align multiple sequences without guide tree (Kruspe and Stadler, 2007). Neighbor-net is the network analogue of the neighbor-joining (NJ) method (Saitou and Nei, 1987). First, a circular ordering of the taxa is computed heuristically, and then a non-negative least squares (NNLS) procedure is applied to find optimal weights for all splits that agree with that circular ordering. The heuristic consists of joining two clusters, using the same criterion as neighbor-joining, and then choosing one taxon from each cluster such that those two taxa will be adjacent in the final circular ordering. The first part of this agglomeration defines a phylogenetic tree on all taxa (Levy and Pachter, 2011) and the second part chooses a circular ordering that agrees with all splits of that tree. It follows from Semple and Steel (2004) that there are 2^*n−*3^ circular orderings with that property for *n* taxa.

Here we present Lpnet, a variant of Neighbor-net that does not apply the second heuristic step of the agglomeration. Instead, we only construct a phylogenetic tree heuristically. Then we use integer linear programming to find an optimal circular ordering, and finally we use the same NNLS procedure as Neighbor-net to assign split weights. We have run experiments to compare Lpnet and Neighbor-net, using random distances, simulated data sets, and a real data set, and we observe that Lpnet tends to approximate the input distance clearly better than Neighbor-net.

In addition to the improved performance compared to Neighbor-net, the strategy to search the possible circular orderings that agree with a binary tree gives rise to a new functionality. A tree can be input by the user, and then Lpnet augments it into a circular split system. This can have many applications, for example if one is confident that all splits of a tree are correct and wants to avoid losing them due to noise in the distance matrix.

Since the integer linear programming problem in Lpnet uses a quadratic number of variables and a cubic number of constraints, Lpnet is computationally more demanding than Neighbor-net. Our implementation in R is feasible for up to 94 taxa and usually needs about one minute on a standard computer for 80 taxa. For larger taxa sets, we provide a top-down heuristic to compute circular ordering from the tree. The method is available at https://github.com/yukimayuli-gmz/lpnet.

### New Approach

As NJ and Neighbor-net, Lpnet uses a matrix with pairwise distances between the taxa as its input. However, all of these methods can be interpreted as quartet-based in the following sense: A quartet *ab*|*cd* consists of two pairs {*a,b*} and {*c,d*} for four different taxa *a,b,c,d*. A weight of *ab*|*cd* can be defined for every quartet, based on the pairwise distances between taxa in {*a,b,c,d*} and quantifies the support for separating *a* and *b* from *c* and *d*. Then all objective functions and selection criteria of the agglomerative part of the considered distance-based methods can be written as functions of the weights of some quartets. This has been noted by Mihaescu *et al*. (2009) and Levy and Pachter (2011), and we will give details in the Materials and Methods section, but now we will only use the quartet interpretation to describe the differences between Lpnet and Neighbor-net. The Lpnet algorithm can be divided into three steps:

1. Construct a phylogenetic tree from the input distance matrix. We use several methods to do this construction, including NJ.
2. Use Integer Linear Programming to find a circular ordering which is consistent with the tree to maximize the sum of all quartet weights contained in the circular ordering. If the number of taxa exceeds 94, compute an ordering heuristically.
3. Estimate split weights from the circular ordering by using non-negative least squares.

The tree construction and the split weights estimation are similar to what NJ, Neighbor-net and other tree or network algorithms do. We only sketch the part using Linear Programming in this section, because this step is the main distinction between Lpnet and Neighbor-net.

Neighbor-net combines the agglomeration of bottom-up clustering with a second selection step that essentially defines an ordering of the taxa in the newly added cluster. Every cluster can be considered a path with its taxa as vertices, and the path will be an interval in the final circular ordering. When two clusters are merged, one end vertex from each path is selected, and those vertices are connected to a new path for the union of the clusters. The selection maximizes a weighted sum of all quartets that are supported by every circular ordering containing the merged path but not by every ordering containing the two short paths. Crucially, the decision about the ordering of the new cluster is made based on the information available at the time of merging.

For Lpnet, we choose to delay the ordering of the clusters until the whole tree is known. We do so by solving an integer linear programming problem that finds a circular ordering of all taxa that maximizes the sum of the weights of its supported quartets among all orderings that agree with all splits of the tree.

### Using Linear Programming to maximize quartet weights

When a phylogenetic tree is drawn in the plane, this embedding defines a circular ordering of the taxa which can be observed by traversing the tree such that every edge is visited once in each direction (see Figure 1). The circular orderings associated with a given tree have been studied by Semple and Steel (2004) who showed that, for a binary tree with *n* taxa, there are exactly 2^*n−*3^ orderings. Starting with an embedding, every possible ordering can be obtained by flipping an arbitrary subset of the interior edges.

**FIG. 1.**
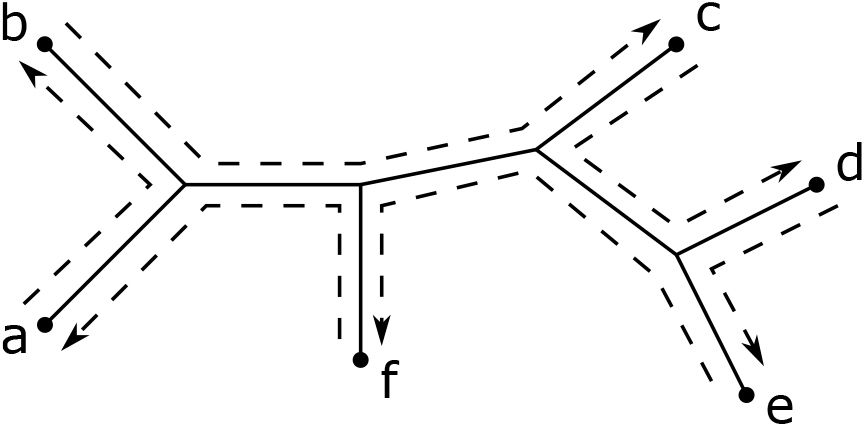
A phylogenetic tree is drawn in the plane. Traversing its edges gives the circular ordering (a,b,c,d,e,f).

In terms of supported quartets, for every four taxa *a,b,c,d*, there is one quartet, say *ab*|*cd*, that is displayed by the tree in the sense that there is at least one split separating *a,b* from *c,d*. Every circular ordering that agrees with the tree will support *ab*|*cd* and one of the other two quartets *ac*|*bd, ad*|*bc*. If the ordering of an initial embedding of the tree supports *ad*|*bc*, then another ordering will support *ad*|*bc*, if and only if the number of flipped edges separating *a,b* from *c,d* is even. We search all orderings that agree with a tree by starting with an initial embedding and introducing a binary variable *X*_*i,j*_ for every pair of interior vertices which will be set to 1, if an odd number of edges on the path between *i* and *j* is flipped, and to 0 else. We give an example of a 5-taxa tree with all four possible circular orderings in Figure 2, and we list the values of the variables. We introduce linear constraints that make sure that the allowed assignments to the variables correspond precisely to the circular orderings that agree with the tree. Then the sum of the weights of all supported quartets that are not displayed by the tree is a linear function of the variables, and we can globally maximize this objective function using binary linear programming.

**FIG. 2.**
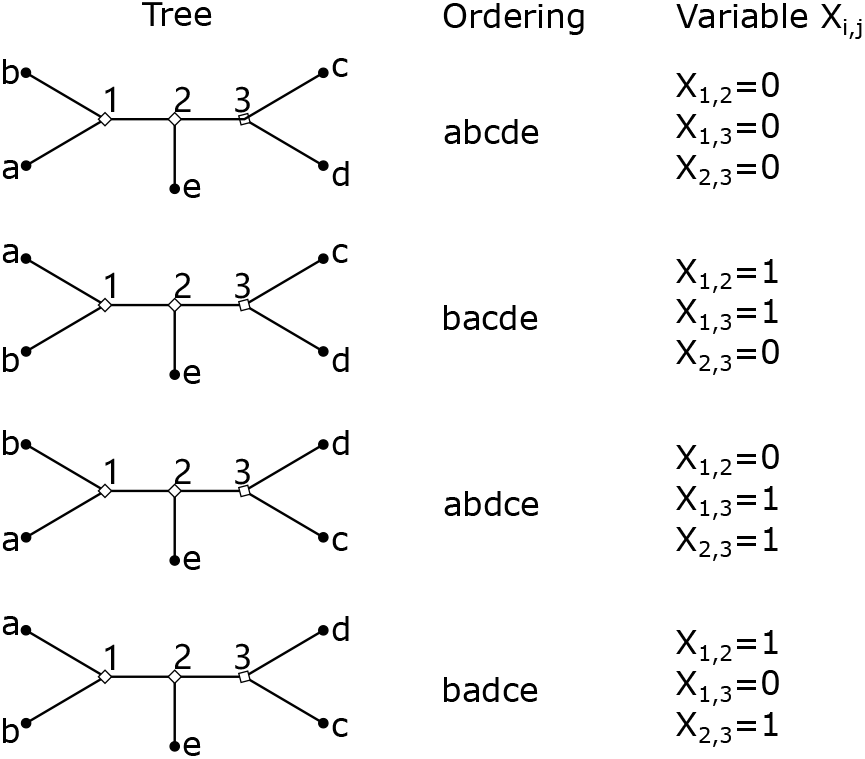
A 5-taxa tree and the correspondence between circular orderings and variables *X*_*i,j*_.

### Maximizing quartet weights heuristically

Even if an exact solution to maximize the quartet weights is not feasible, it has some advantages to first complete the tree construction and then 4 compute a circular ordering. We include a top-down heuristic which iteratively fixes the variables *X*_*i,j*_ for edges *ij*. In contrast to Neighbor-net, it uses all sets of 4 taxa. It also can choose between several candidate variables to be fixed first, and it revises its decisions in a post-processing step.

## Results

In this section, we mainly compare the performance of Neighbor-net and Lpnet for different input distances. In order to evaluate different networks from the same input, we use the LSFit, which is also available in SplitsTree4 (Huson and Bryant, 2006). It takes the sum of the squares of the differences between input and output distances, and rescales this number such that a perfect match yields an LSFit of 100 while setting all distances to zero yields 0. Before the split weights are computed, Neighbor-net and Lpnet both try to maximize the sum of the weights of the quartets that agree with the chosen circular ordering. Therefore, we use this sum to evaluate the algorithms at that state and refer to it as *sum of quartets*. Within Lpnet, the result may depend on the choice of the tree reconstruction method. We use Neighbor-joining and its common variants BioNJ (Gascuel, 1997a) and UNJ (Gascuel, 1997b), as well as two methods that try to mimic the internal tree building of Neighbor-net. We call them *NNet tree* and *symmetric NNet tree* (see the Materials and Methods section for details).

We run examples on four different kinds of input distances: First we present a simple artificial example with only seven taxa and two reticulations that shows how the early ordering of clusters by Neighbor-net can cause problems. The second example uses random distances between taxa and represents the general approximation problem of an input metric by a circular split system, without any phylogenetic signal. The third example uses simulated sequences from random trees. The last example is a published data set of *Viburnum* plants which contains a hypothesized hybrid and has been suggested as a benchmark for phylogenetic networks methods.

### An artificial example

We start with an artificial example that demonstrates the disadvantage of ordering clusters locally. As visualized in Figure 3, we assume that seven taxa mainly evolved under a clocklike tree, with the exception of two gene transfer events *I*_1_ and *I*_2_. We assume that 10% (for *I*_1_) respectively 20% (for *I*_2_) of the genome of the reticulation vertices are independently replaced along the reticulation arrows. This means that the genome consists of four parts representing sequences affected by one or both or none of the reticulations. Each part follows its own tree, and the observed distance is a convex combination of those four tree distances where the coefficients are the fractions of the genome that follow the trees. This distance corresponds to ten non-trivial splits which do not fit on any circular ordering, and seven trivial splits. The non-trivial splits with the weights and the contributing trees are listed in Table 1, and the distance matrix representing the network is given in Table 2.

**FIG. 3.**
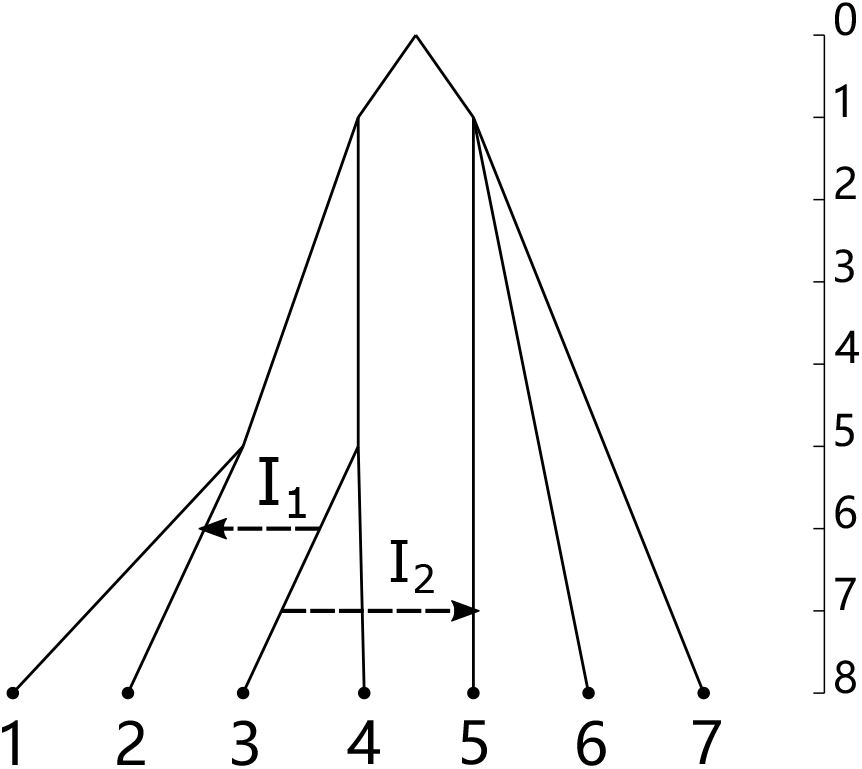
A tree-like phylogentic network with two reticulations.

**Table 1.** Nontrival splits and positive weights for the original phylogenetic tree and the three trees representing transfer events, the mixture of the input trees and for Neighbor-net and Lpnet.

| original tree (72%) | | $TI_1$ (8%) | | $TI_2$ (18%) | | $TI_{12}$ (2%) | | split | Tree mixture weight | NNet weight | Lpnet weight |
| --- | --- | --- | --- | --- | --- | --- | --- | --- | --- | --- | --- |
| split | weight | split | weight | split | weight | split | weight |  |  |  |  |
| 12 | 4 |  |  | 12 | 4 |  |  | 12\|34567 | 3.6 | 3.6 | 3.6 |
| 34 | 4 |  |  |  |  |  |  | 12567\|34 | 2.88 | 2.65 | 2.8375 |
| 567 | 2 | 567 | 2 |  |  |  |  | 1234\|567 | 1.6 | 1.6 | 1.6 |
|  |  | 23 | 1 |  |  |  |  | 14567\|23 | 0.08 | 6.215 e-16 |  |
|  |  | 234 | 4 |  |  |  |  | 1567\|234 | 0.32 | 0.35 | 0.35 |
|  |  |  |  | 35 | 2 | 35 | 1 | 12467\|35 | 0.38 |  | 0.375 |
|  |  |  |  | 345 | 4 |  |  | 1267\|345 | 0.72 | 0.9 | 0.7125 |
|  |  |  |  | 67 | 2 | 67 | 2 | 12345\|67 | 0.4 | 0.4 | 0.4 |
|  |  |  |  |  |  | 167 | 4 | 167\|2345 | 0.08 | 0.1 | 0.1 |
|  |  |  |  |  |  | 235 | 1 | 1467\|235 | 0.02 |  |  |
|  |  |  |  |  |  |  |  | 12347\|56 | 0 |  | 3.73 e-16 |

NOTE.—The contributing trees and their percentage of the genome are listed in the header. Every row corresponds to a non-trivial split. For every contributing tree, we represent its splits by the smaller split halves and list the split weights. The correct weight for every non-trivial split throughout the genome is given in the *Tree mixture weight* column.

**Table 2.** The distance matrix for our artificial example.

|  | 1 | 2 | 3 | 4 | 5 | 6 |
| --- | --- | --- | --- | --- | --- | --- |
| 2 | 6.8 |  |  |  |  |  |
| 3 | 14 | 13 |  |  |  |  |
| 4 | 14 | 13.2 | 6 |  |  |  |
| 5 | 15.6 | 15.4 | 13.2 | 14 |  |  |
| 6 | 16 | 16 | 16 | 16 | 14.4 |  |
| 7 | 16 | 16 | 16 | 16 | 14.4 | 14 |

When Neighbor-net is applied to this distance, it correctly identifies the clusters {1,2} and {3,4} first and then decides to join those clusters. The second selection criterion of Neighbor-net chooses to make 2 and 3 neighbours in the circular ordering, because it relies on quartets that have at most one taxon from {5,6,7}. This decision based on local information makes it impossible to later include the cluster {3,5}, which corresponds to a much stronger split than {2,3}. Lpnet first correctly finds the whole main tree and then chooses a circular ordering that allows all true splits with a weight higher than 0.1.

As can be seen from Table 1, the weights of the correct splits in Lpnet tend to be clearly closer to the true weight than Neighbor-net, and even 23|14567, the only true split that is allowed by the Neighbor-net ordering and not by the Lpnet one, finally gets a negligible weight in the Neighbor-net. Both methods achieve a very high LSFit (Neighbor-net 99.99445 and Lpnet 99.99965), but the gap to 100 is still more than fifteen times greater for Neighbor-net than for Lpnet.

### Random distances

We assign a random number *r*(*x,y*) between 0 and 1 from the uniform distribution to every pair *x,y* of taxa. In order to guarantee the triangle inequality, we define the distance between *x* and *y* to be *d*(*x,y*) = *r*(*x,y*)+*c*, where *c* is the maximum of *r*(*x,z*)*−r*(*x,y*)*−r*(*y,z*) over all three taxa *x,y,z*.

We generate 10000 random distance matrices with 30 taxa. Then we compare Lpnet using different methods to construct phylogenetic trees, with Neighbor-net. We use SplitsTree4 (Huson and Bryant, 2006) with default setting to get the Neighbor-net result. Then we compare the sum of quartets and the LSFit value for Lpnet and Neighbor-net (Table 3). We observe that for all five tree building methods, for the sum of quartets and LSFit, the Lpnet algorithm clearly tends to get better scores than Neighbor-net. While Lpnet achieves a higher LSFit for roughly 80% of the input metrics, this fraction is more than 98% for the sum of quartets. Comparing the Lpnets using different tree building methods, we find that we often get the same circular ordering. Nevertheless, Table 4 shows that UNJ performs significantly better than the other methods. Our heuristic, based on the UNJ tree, is a clear improvement compared to Neighbor-net, but it produces worse results than Integer Linear Programming.

**Table 3.** Comparison of Lpnets (five different methods of tree construction) and heuristic method with Neighbor-net in terms of the LSFit value and the sum of quartets, for 10000 random distances.

|  | Same circular ordering<br>as Neighbor-net | LSFit |  | sum of quartets |  |
| --- | --- | --- | --- | --- | --- |
|  |  | Lpnet > NNet | Lpnet < NNet | Lpnet > NNet | Lpnet < NNet |
| Neighbor-joining | 0 | 7782 | 2218 | 9828 | 172 |
| Symmetric NNet tree | 0 | 8259 | 1741 | 9974 | 26 |
| NNet tree | 0 | 8272 | 1728 | 9967 | 33 |
| UNJ | 0 | 8251 | 1749 | 9977 | 23 |
| BioNJ | 0 | 7962 | 2038 | 9903 | 197 |
| Heuristic method | 0 | 7200 | 2800 | 9872 | 128 |

**Table 4.** Comparison of Lpnets using UNJ with Lpnets using four other tree reconstruction methods and heuristic method for 10000 random distances.

|  | Same circular ordering<br>as UNJ | LSFit |  | sum of quartets |  |
| --- | --- | --- | --- | --- | --- |
|  |  | > UNJ | < UNJ | > UNJ | < UNJ |
| Neighbor-joining | 1107 | 3989 | 4904 | 3370 | 5523 |
| Symmetric NNet tree | 5427 | 2273 | 2300 | 2038 | 2535 |
| NNet tree | 3909 | 2981 | 3110 | 2556 | 3535 |
| BioNJ | 2138 | 3759 | 4103 | 3621 | 4241 |
| Heuristic method | 2807 | 1951 | 5242 | 0 | 7193 |

Since there is no phylogenetic signal in the random distances, there are often many different circular orderings with similar residuals. The high percentage of the examples where Lpnet achieves a lower residue than Neighbor-net shows that the strategy to construct a binary tree first and choose a circular ordering afterwards improves the distance approximation, even if there is no correct underlying tree or network.

### Simulated sequences

We randomly generate a tree for 30 taxa by using the function ‘sim.taxa’ from the R package TreeSimGM (Hagen and Stadler, 2018). We let the parameter ‘waiting time until speciation’ for ‘sim.taxa’ be exponentially distributed with rate parameter *λ* = 1.2, and then normalize such that the longest pairwise distance is one. Then we use the software Dawg (Cartwright, 2005) to simulate DNA sequences of length 10000bp from the random tree under the Jukes-Cantor model. Finally, we use SplitsTree4 to compute Jukes-Cantor distances with default settings. We repeat this process 10000 times and compare Lpnet and Neighbor-net in the same way as for the random metrics. Table 5 shows the result of comparing the LSFit value and the sum of all quartets between Lpnet and Neighbor-net. We see that for all five tree construction methods, the advantage of Lpnet compared to Neighbor-net increases. The sum of quartets is now always higher and the LSFit better for almost 95% of the data sets when we use Lpnet. The input data sets for this experiment can be interpreted as a tree metric plus some random noise, and the results show that in this situation the strategy of Lpnet to complete the tree reconstruction before embedding the tree pays off. The various tree building methods yield the same circular ordering more often than for random metrics, but again UNJ achieves better scores than the other variants of NJ (see Table 6). In order to compare the performance of the methods for single data sets, we plot the difference of the logarithms of the residuals of Neighbor-net and Lpnet in increasing order in Figure 4. The residual of a network is defined to be the sum of the squares of the differences between the pairwise input distances and the distances induced by the network. The median of this quantity is 0.093, thus for half of the data sets the Neighbor-net residual is by more than 9.7% higher than the Lpnet residual.

**Table 5.** Comparison of Lpnets (five different methods of tree construction) and heuristic method with Neighbor-net in terms of the LSFit value and the sum of quartets, for 10000 simulated sequence alignments.

|  | Same circular ordering<br>as Neighbor-net | LSFit |  | sum of quartets |  |
| --- | --- | --- | --- | --- | --- |
|  |  | Lpnet > NNet | Lpnet < NNet | Lpnet > NNet | Lpnet < NNet |
| Neighbor-joining | 0 | 9406 | 594 | 9997 | 3 |
| Symmetric NNet tree | 0 | 9412 | 588 | 9998 | 2 |
| NNet tree | 0 | 9407 | 593 | 9997 | 3 |
| UNJ | 0 | 9409 | 591 | 9997 | 3 |
| BioNJ | 0 | 9399 | 601 | 9997 | 3 |
| Heuristic method | 0 | 8608 | 1392 | 9728 | 272 |

**Table 6.** Comparison of Lpnets using UNJ with Lpnets using four other tree reconstruction methods and heuristic method for 10000 simulated sequence alignments.

|  | Same circular ordering<br>as UNJ | LSFit |  | sum of quartets |  |
| --- | --- | --- | --- | --- | --- |
|  |  | > UNJ | < UNJ | > UNJ | < UNJ |
| Neighbor-joining | 9276 | 327 | 384 | 266 | 445 |
| Symmetric NNet tree | 9469 | 258 | 272 | 212 | 317 |
| NNet tree | 9466 | 256 | 276 | 210 | 322 |
| BioNJ | 9336 | 307 | 356 | 241 | 422 |
| Heuristic method | 3280 | 1807 | 4897 | 0 | 6704 |

**FIG. 4.**
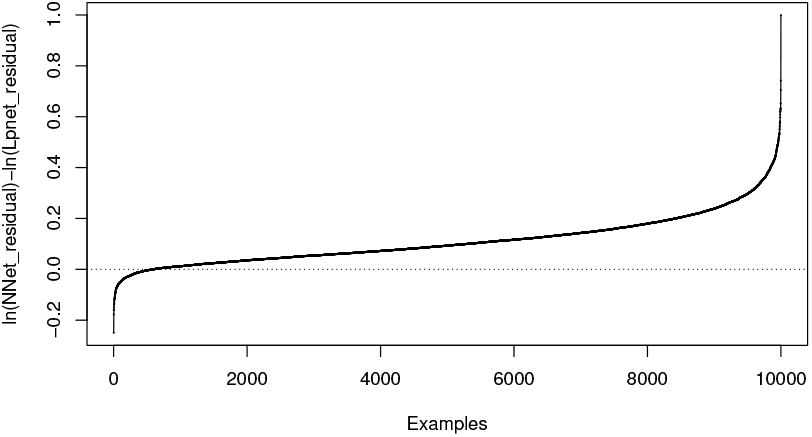
The differences between the logarithms of the residuals of Nnet and Lpnet for the simulated data sets.

### Analysis of a published data set

As an example of an analysis of a real data set, we choose a study of the genus *Viburnum* of flowering plants (Donoghue *et al*., 2004). The raw data are chloroplast trnK intron and nuclear ribosomal ITS DNA sequences from 43 species of *Viburnum* and 2 species of *Sambucus*. We use the uncorrected P distance from the combined sequence alignment to compute the Neighbor-net (Fig 5) and the Lpnet (Fig 6). Following the previous results, we chose UNJ as the tree reconstruction method for Lpnet.

**FIG. 5.**
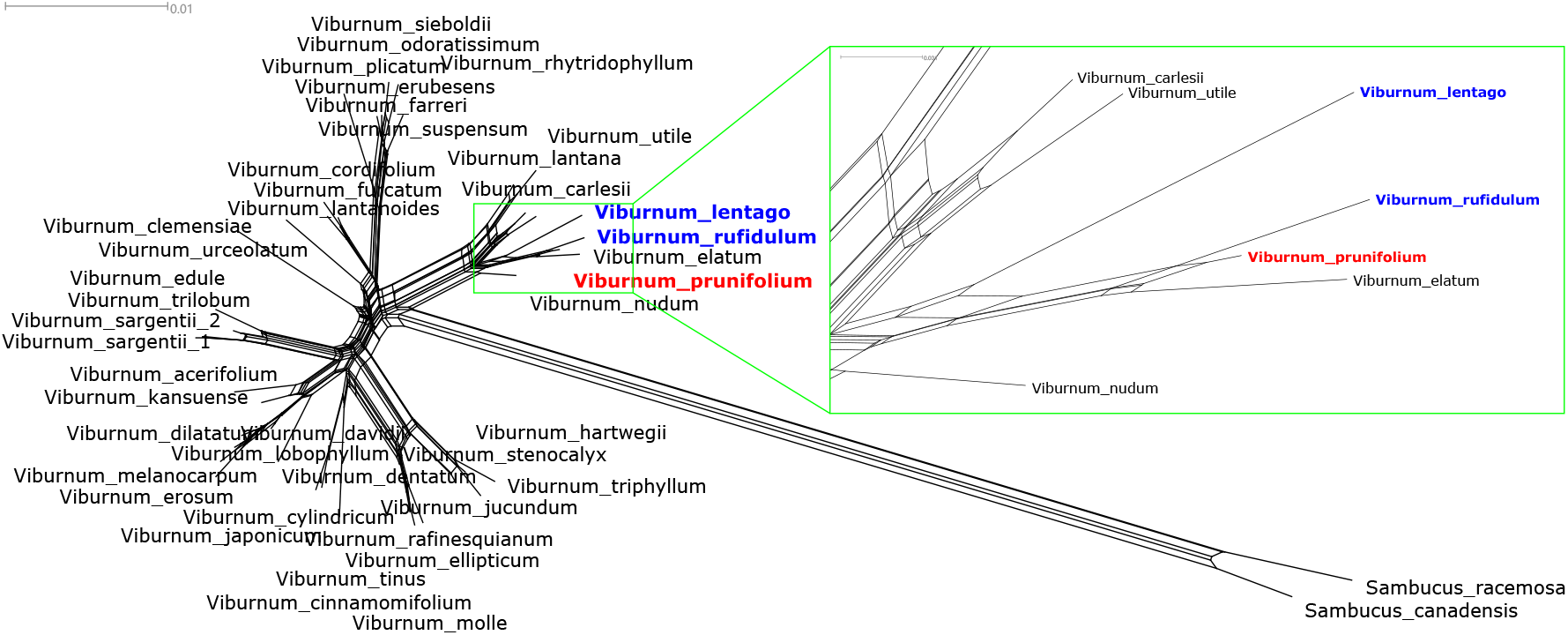
Neighbor-net. Viburnum_prunifolium (red) is the hybrid between Viburnum_rufidulum and Viburnum _lentago (blue).

**FIG. 6.**
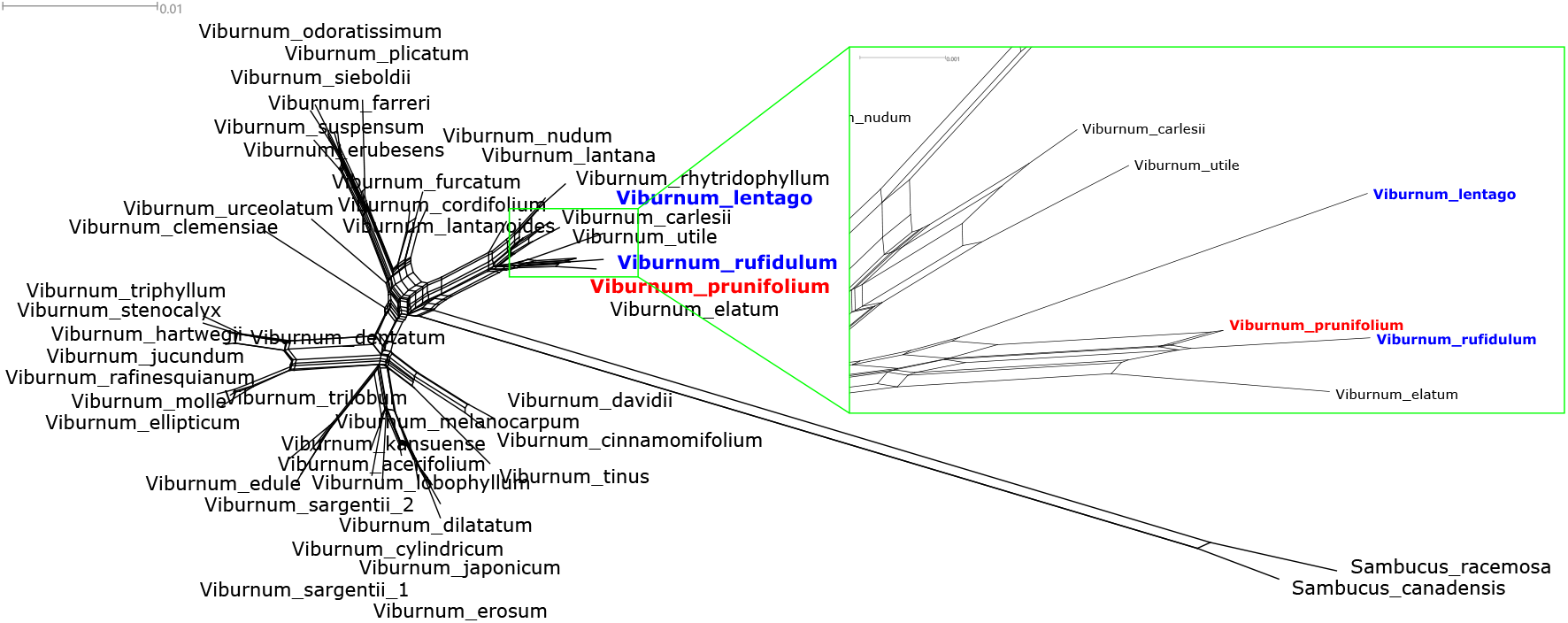
Lpnet. Viburnum_prunifolium (red) is the hybrid between Viburnum_rufidulum and Viburnum lentago (blue).

This data set has been proposed as an example for testing phylogenetic networks method by an influential but now inactive blog (phylonetworks.blogspot.com/p/datasets.html). We focus on the position of *V. prunifolium* which was already hypothesized to be a hybrid between *V. lentago* and *V. rufidulum* in 1956 (Brumbaugh and Guard, 1956). From the differences between the trees obtained by analyzing the two loci separately, Donoghue *et al*. (2004) conclude that their data set supports that hypothesis. In Figs 5 (NNet) and 6 (Lpnet), we highlight *V. prunifolium* in red, and *V. lentago* and *V. rufidulum* in blue. We observe that in the Neighbor-net, *V. prunifolium* is not placed between *V. lentago* and *V. rufidulum*, and there is no split separating *V. prunifolium* and *V. lentago* from all other taxa and no split separating *V. prunifolium* and *V. rufidulum* from all other taxa. In the Lpnet, *V. prunifolium* is between *V. lentago* and *V. rufidulum*, and the 2-splits grouping together only *V. prunifolium* and *V. lentago* (with medium weight) and grouping together *V. prunifolium* and *V. rufidulum* (with low weight) are both present.

From Donoghue’s study (Donoghue et al., 2004), we observe that there is a cluster containing only *V. prunifolium* and *V. lentago* in the phylogenetic tree for the trnK alignment and an unresolved cluster containing *V. prunifolium, V. rufidulum*, and *V. elatum* for the ITS alignment. The latter split is strong and can be observed in both networks, while the former split has medium weight in the Lpnet and conflicts with the circular ordering of the Neighbor-net. This indicates that the distance matrix is indeed better represented by the Lpnet, which is also confirmed by the better LSFit value (Lpnet: 99.90413, Nnet: 99.884).

## Discussion

Seventeen years after the release of Neighbor-net, our new variant Lpnet provides an alternative that approximates the input distance better for the clear majority of the data sets we have tried. The main disadvantage of Lpnet is that it is slower and needs more memory, but most data sets that have been analyzed by Neighbor-net have less than 80 taxa and can therefore be handled by Lpnet as well.

The main application of split graphs and in particular Neighbor-nets is to find the main signals in an early stage of a data analysis (Huson and Bryant, 2006). In practice, a Neighbor-net often contains a few strong splits and many tiny ones which are usually interpreted as irrelevant noise. We expect that the clear signals will often be detected by both methods, while differences between the minor splits will cause a slightly higher LSFit value for Lpnet. In such cases it does not matter much which method is used.

However, our realistic artificial example demonstrates that there can be significant differences. If reticulations are supported by a data set, then the goal has to be to reconstruct an explicit phylogenetic network as shown in Figure 3. While it is generally hard to guess that network from a splits graph, this task would be easier for the Lpnet than for the Neighbor-net. We therefore anticipate that Lpnet will be useful for interpreting real data sets in the future.

We provide five different algorithms to construct phylogenetic trees. All of them turn out to yield the best LSFit occasionally, and we have no strong preference. In the average, UNJ performed best for our data sets, and it seems most consistent with the general approach taken by Lpnet that treats all pairs of taxa and all quartets equal. Therefore, we select UNJ as the default method, but we recommend to try other methods as well. Other tree building methods like minimum evolution or even not distance-based methods like maximum likelihood might be worth trying, and any binary tree can be input to Lpnet. It is generally interesting to see whether all splits of a reasonable tree will have positive weights in the Lpnet, and to compare the weights of the strongest other splits with the weights of the conflicting tree splits. However, users should be aware that we only have a consistency proof for the neighbor-joining variants that we provide.

Even though we replaced a heuristic part of Neighbor-net by an exact algorithm, Lpnet is still a heuristic method. It relies on a heuristic tree construction, and it optimizes the sum of quartet weights, while the final score function for a weighted split system is the LSfit value. The weights of the supported quartets indicate but do not guarantee that the distance can be approximated well by the allowed splits, and the discrepancy causes almost all cases where the LSFit of Lpnet is worse than Neighbor-net. It would be desirable to have a method that directly optimizes the least squares fit, but this would not allow any agglomerative construction and we are not aware of any such algorithm other than trying every possible ordering.

QNet (Grünewald *et al*., 2007) is the quartet analogue of Neighbor-net. It uses quartet weights directly obtained from the raw data instead of distances to reconstruct a weighted circular split system. The strategy of Lpnet to first construct a tree and then use linear programming to get a circular ordering, can also be applied to modify QNet, and the approximation is expected to improve.

Another direction of potential future work is to change the score that is used to compute the split weights. Using non-negative least squares is an intuitive choice, especially if there is no information about the distribution of the error (the difference between the observed and the correct distances). If the input distances are computed from a sequence alignment, it is plausible that short distances have lower variance than long ones, and weighted least squares have been used for estimating the edge lengths of a tree for a long time (Fitch and Margoliash, 1967). A non-negative version might be useful for circular split systems. If a model is available that assigns a likelihood of the observed distance matrix to every potential underlying true distance, then maximizing that likelihood is an alternative to least squares optimization. For all these variants, we expect it to be advantageous to first construct a binary tree and then embed it into a circular ordering.

## Materials and Methods

### Compatible splits and circular split systems

A split *S* divides a set *X* of taxa to two nonempty parts *A* and *B* and is denoted *S* = *A*|*B*. Two splits *A*_1_|*B*_1_ and *A*_2_|*B*_2_ are *compatible*, if at least one of the intersections *A*_1_ *∩A*_2_, *A*_1_ *∩B*_2_, *B*_1_ *∩A*_2_, *B*_1_ *∩ B*_2_, is empty. For a phylogenetic tree where the leaf nodes are labeled by the set *X* of taxa, every branch of the tree represents a split of *X* and all splits are compatible with each other (Semple and Steel, 2003).

Three splits *S*_1_ = *A*_1_|*B*_1_, *S*_2_ = *A*_2_|*B*_2_ and *S*_3_ = *A*_3_|*B*_3_ are *weakly compatible*, if there are no four taxa {*t,u,v,w*} with {*t,u} ∈ A*_1_, {*v,w} ∈ B*_1_, {*t,v} ∈ A*_2_, {*u,w} ∈ B*_2_, {*t,w} ∈ A*_3_, {*u,v} ∈ B*_3_ (Bandelt and Dress, 1992). A *circular ordering* of a set *X* = {*x*_1_,…,*x*_*n*_} of *n ≥* 3 taxa can be obtained by labelling all vertices of an *n*-gon by the taxa (see Fig 7). A split *A*|*B agrees* with a circular ordering, if both *A* and *B* label consecutive paths on the circle. A circular ordering is defined by the permutation of *X* that we get by starting at an arbitrary taxon and then following all vertices of the cycle in a clocklike or anti-clocklike fashion. Therefore, there are 2*n* different permutations associated with the same circular ordering.

**FIG. 7.**
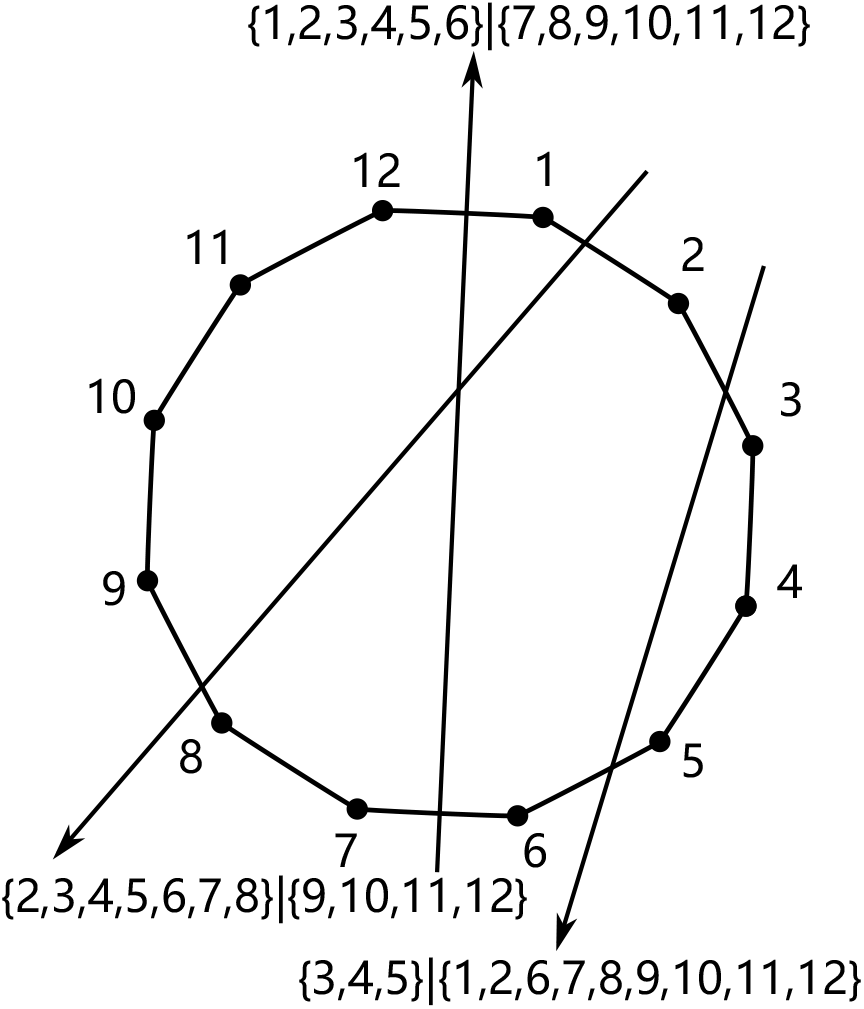
A circular split system and three splits.

A *circular split system* of *X* is a set of splits of *X* such that all splits agree with a single circular ordering. It has long been known that circular split systems are weakly compatible and have up to ^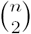^ splits (Bandelt and Dress, 1992). Further, compatible split systems are circular, thus an unrooted phylogenetic tree can be considered a circular split system. When non-negative weights are assigned to the splits, this *weighted circular split system* can be visualized by a planar *splits graph*, a network where the taxa are embedded as vertices, and every split *A*|*B* of weight *w* corresponds to a set of parallel edges which all have length *w*. Further, removing all those edges decomposes the network into two connected components containing *A* and *B*, respectively (Dress and Huson, 2004). Splits graphs are commonly referred to as implicit phylogenetic networks, and SplitsTree4 can be used to draw such a network from an input weighted circular split system (Huson and Bryant, 2006).

Every weighted split system induces a pairwise distance on the taxa where the distance between two taxa *x* and *y* is the sum of the weights of all splits that separate *x* from *y*.

### Quartets and weights

As mentioned before, Neighbor-joining and its variants can be considered a quartet method (Mihaescu *et al*., 2009). For a distance *d*, and four taxa *u,v,x,y*, we define the weight of the quartet *uv*|*xy* by *w*(*uv*|*xy*) = *d*(*u,x*)+*d*(*u,y*)+ *d*(*v,x*)+*d*(*v,y*)*−*2*d*(*u,v*)*−*2*d*(*x,y*). If *d* is induced by a weighted split system, then *w*(*uv*|*xy*) equals four times the sum of the weights of all splits that separate *u,v* from *x,y* minus two times the sum of the weights of all splits that support a conflicting quartet. Therefore, the weight of a quartet quantifies the support for separating its pairs. The support for a tree (without edge lengths) or a circular ordering can be quantified by summing up the weights of all supported quartets, and this number is maximized by a correct tree or ordering, if the input distance is induced by a tree or a circular split system, respectively.

### Distance reduction and the order dependence of Neighbor-net

The first step of neighbor-joining and all other agglomerative algorithms discussed here is to identify two taxa *u,v*, such that the sum of all quartet weights *w*(*uv*|*xy*) is maximized. Then the cluster {*u,v*} is considered a single taxon, where the distances between this new taxon and another taxon *x* is is based on the distances between *u, v*, and *x*. This process is reiterated until there are only three taxa left, and the two steps are called the *selection* and the *reduction* step. It was pointed out by Bryant (2005) that, in order to reconstruct trees correctly from their induced metric, the selection step is unique, while the reduction step can give different weights to the two joined clusters. Neighbor-joining always gives the same weight to both clusters, while UNJ uses weights that are proportional to the cluster size, and BioNJ tries to minimize the variance of the reduced distances.

Neighbor-net does not reduce the distance for clusters of size two, because the distinction helps to compute a circular ordering agglomeratively. Instead, it reduces three taxa to two, whenever a cluster of size two is merged with another cluster. If both clusters have size two, this reduction step has to be performed twice. This distinguishes the taxon that is not included in the first reduction from the other three. For default parameters, the taxa that are first reduced each receive 2*/*9 and the remaining taxon 1*/*3 of the total weight of the new cluster. Since the choice which three taxa are reduced first is not determined by the input distances, the output of Neighbor-net can sometimes depend on the input order of the taxa, even if no ties occur. It therefore seems reasonable to give equal weights to all four clusters. We have implemented this variant of the Neighbor-net tree construction in Lpnet and refer to it as *symmetric NNet tree*. In addition, we use *NNet tree* to mimic the original weighting by randomly assigning 1*/*3 to one of the two candidate taxa that might receive that weight from the Neighbor-net algorithm.

### The Lpnet algorithm

As we have said in the New Approaches section, the Lpnet algorithm uses a distance matrix as its input. First it constructs a phylogenetic tree from the distances, then it uses Linear Programming to find a circular ordering which maximizes the sum of all quartet weights consistent with the circular ordering. Finally, it computes weights for all splits that agree with the circular ordering such that the LSFit is maximized.

#### Constructing a tree

Neighbor-net constructs a circular ordering agglomeratively. The process is the same as for QNet (Grünewald *et al*., 2007), where it is described as adding edges to a graph whose vertices are the taxa such that every component is a path. It starts with the graph without edges and joins two paths in every iteration, until a single path is obtained which then defines a circular ordering. All taxa sets of the paths form a compatible cluster system and therefore define a binary tree. The difference to methods that only construct a tree is that after selecting two paths to be merged, there are up to four pairs of taxa that can be connected to get a single path. One such pair is chosen by the second selection criterion of Neighbor-net which maximizes the sum of the weights of all quartets that are supported by all circular orderings containing the new path.

As indicated in the previous subsection, the second selection step has some influence on the tree construction. It was already suggested by Levy and Pachter (2011) to change the distance reduction of Neighbor-net such that all splits of the neighbor-joining tree are always supported by the output ordering. One main idea of Lpnet is to skip the second selection altogether and choose a circular ordering after the tree construction. In order to observe how much of the performance difference between Lpnet and Neighbor-net is caused by this new strategy, we include the NNet tree which stays as close as possible to Neighbor-net. Noting the undesired order dependence of Neighbor-net, we also include the symmetric NNet tree.

The other tree reconstruction methods implemented by Lpnet are neighbor-joining and its variants, UNJ and BioNJ. It is also possible to input a user-defined tree, and Lpnet will compute a circular split system where all splits of the input tree agree with the circular ordering.

#### Using Linear Programming to maximize quartet weights

Given a binary unrooted tree *T* with *n* taxa and pairwise distances, we want to find a circular ordering that agrees with all splits of *T*, such that the sum of the weights of all supported quartets is maximized. Given an initial ordering that agrees with *T*, we can obtain another such ordering by choosing a non-trivial split *A*|*B* and reversing the order of *A*. Note that reversing the order of *B* yields the same circular ordering as reversing *A*, and reversing both yields the initial circular ordering. This process can be interpreted as flipping the edge that separates *A* from *B*, and it follows from Semple and Steel (2004) that all allowed circular orderings can be obtained by a sequence of edge flips. Moreover, the final circular ordering does not depend on the order in which the edges are flipped, flipping an edge twice yields the same circular ordering as never flipping it, and flipping two different sets of edges always results in different circular orderings. Therefore, there are 2^*n−*3^ different circular orderings that agree with *T*, and they are in one-to-one correspondence with the subsets of the interior edges of *T*.

For four taxa *i*_1_,*i*_2_,*j*_1_,*j*_2_, assume that the quartet *i*_1_*i*_2_|*j*_1_*j*_2_ is displayed by *T*. Then all edges corresponding to splits that separate *i*_1_ and *i*_2_ from *j*_1_ and *j*_2_ form a path in *T*. One of the end vertices of that path, say *i*, is on the path from *i*_1_ to *i*_2_ and the other one, say *j*, is on the path from *j*_1_ to *j*_2_. If the initial circular ordering supports the quartet *i*_2_*j*_1_|*j*_2_*i*_1_, then another circular ordering also supports that quartet, if the number of flipped edges on the path from *i* to *j* is even, else it supports *i*_1_*j*_1_|*i*_2_*j*_2_. Let *I*_1_,*I*_2_,*J*_1_,*J*_2_ be the sets of taxa that are in the same component of the graph obtained from *T* by removing *i* and *j* as *i*_1_,*i*_2_,*j*_1_,*j*_2_, respectively. Then we define

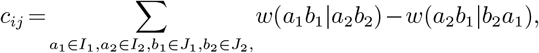

so *c*_*ij*_ quantifies how much we would like to flip an odd number of edges on the path from *i* to *j*. Introducing binary variables, *X*_*ij*_ for every pair {*i,j*} of interior vertices of *T*, we need to find a circular ordering such that *X*_*ij*_ = 1, if and only if the number of flipped edges on the path from *i* to *j* is odd, and Σ_*i,j*_ *c*_*ij*_*X*_*ij*_ is maximal. It turns out that a (0,1)-assignment to all variables corresponds to a circular ordering, if and only if the following four linear inequalities hold for every three interior vertices *i,j,k* of *T* :

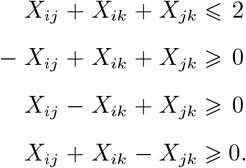

These conditions are necessary, because every edge of the smallest subtree of *T* containing *i,j,k* is contained in exactly two of the three paths between two of those vertices. This means that the sum *X*_*ij*_ +*X*_*ik*_ +*X*_*jk*_ is even, so either all three variables are zero or there are two ones and one zero. To see that the conditions are sufficient, we note that the (0,1)-assignment to all those variables *X*_*ij*_ where *ij* is an edge of *T* already defines a circular ordering. Now there is a single extension of this assignment to all variables such that all conditions hold: Let *i* and *k* be two interior vertices of *T* such that *X*_*ik*_ is unknown while *X*_*ij*_ and *X*_*jk*_ have already been assigned for a vertex *j*. As before, *X*_*ij*_ +*X*_*ik*_ +*X*_*jk*_ must be even, so there is only one allowed assignment for *X*_*ik*_.

In summary, we compute an optimal circular ordering by solving a binary linear programming problem with 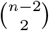 variables and 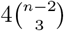 constraints.

#### Maximizing quartet weights heuristically

The agglomeration process of NJ and its variants ends when there are only three clusters left, and for each of those clusters a rooted binary tree has been constructed. This corresponds to a rooted tree where the root has outdegree 3 which we will denote *T*. We use that tree and an initial circular ordering to decide in a top-down fashion which of the edges should be flipped. We assume that there is a subtree *U* containing the root such that for all edges of *U* a decision has been made. Initially, the subtree contains only the root and no edges. For every interior edge of *T* that has exactly one vertex *u* in *U* and the other vertex *v* outside *U*, we compute the average weight of all quartets that are displayed if and only if the edge *uv* is flipped minus the average weight of the quartets that are displayed if and only if *uv* is not flipped. A 4-set {*x*_1_,*x*_2_,*x*_3_,*x*_4_} of taxa contributes to that score, if and only if, for the smallest subtree of *T* connecting *x*_1_,*x*_2_,*x*_3_,*x*_4_, one of the two vertices of degree 3 is *v*, and the other one is in *U*. For the edge *uv* that maximizes the absolute value of this difference, we decide to flip the edge, if the difference is positive, or to not flip it. Then we add the vertex *v* and the edge *uv* to *U* and go to the next iteration. The process stops when *U* contains all interior edges of *T*. After this top-down procedure, we do some post-processing by reversing the decision whenever doing so increases the sum of the quartet weights. In order to guarantee polynomial running time, we stop this process, either when we find no edge that improves the score, or when every edge has been checked *n/*2 times where *n* is the number of taxa.

#### Estimate split weights by non-negative least squares

The last part of Lpnet is to compute weights for all splits that agree with the circular ordering that was constructed in the previous step. Here we follow the same approach as Neighbor-net and use the NNLS algorithm by Lawson and Hanson (1995) to obtain split weights such that the difference between the input distance and the distance induced by the weighted circular split system is minimized. More precisely, we minimize the Euclidian norm of 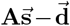 where **A** is the (0,1)-matrix with rows indexed by the pairs of taxa and columns indexed by the allowed splits, and an entry is one, if and only the pair of taxa corresponding to its row is separated by the split corresponding to its column. Further, 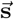 and 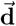 are the column vectors with the non-negative split weights and the input distances, respectively. In order to measure the goodness of distance approximation, we use the least squares fit (LSFit):

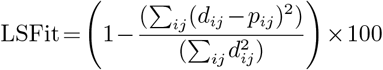

where the pairwise distances induced by the weighted split system are *p*_*ij*_ and the input distances are *d*_*ij*_.

### Consistency of Lpnet

Consistency is an important feature of phylogenetic reconstruction methods. It means that a method does not make mistakes for perfect input. Specifically, a method that reconstructs a circular split system from distances is *consistent*, if it returns the correct weighted split system whenever the input distance is induced by a weighted circular split system. Neighbor-net was shown to be consistent by Bryant *et al*. (2007). A more general proof was given by Levy and Pachter (2011), where all tree construction methods used by Lpnet are included in their definition of neighbor-joining which allows a wide class of weighting schemes for the reduction step. Their result implies that all splits of the tree constructed by Lpnet agree with some circular ordering that agrees with all splits of the underlying split system. It is easy to see that such an ordering will also maximize the sum of all supported quartets. Finally, NNLS will be able to match the input distance exactly, thus Lpnet is consistent for all used tree reconstruction methods.

### Implementation

We provide an R implementation of Lpnet that allows the user to choose one of the five tree reconstruction methods listed above or to input a tree. For the linear programming problem, two solvers are supported: The R version Rglpk of the GNU Linear Programming Kit (http://www.gnu.org/software/glpk) is free and open source, while Gurobi (http://www.gurobi.com) is one of the most powerful commercial mathematical optimization solvers. A free license of Gurobi is available for academic institutions.

In practice, the integer linear programming is the computationally most demanding part of Lpnet. The constraint matrix has *O*(*n*^5^) entries, and the required memory grows equally fast. There is a hard limit of 94 taxa, because the matrix is stored as a vector, and R only allows vectors of length at most 2^31^ *−*1. Using Gurobi, we were able to compute solutions for all input distances with up to 94 taxa that we tried.

We list the size of the constraint matrix and the average CPU time for running our Lpnet function in R using Gurobi with binary linear programming for different numbers of taxa in Table 7. Rglpk is much slower, and the problem was often not feasible for more than 50 taxa on our machine. For 40 taxa, we observed an average CPU time of 13.76 seconds.

**Table 7.** Lpnet. Sizes of the constraint matrix and User times for different number of taxa

| number of taxa | User time | size of constraint matrix |
| --- | --- | --- |
| 20 | 0.0803 sec | 3.81 MB |
| 40 | 1.4167 sec | 180.985 MB |
| 60 | 11.7401 sec | 1.52 GB |
| 80 | 59.8303 sec | 6.809 GB |

Integer linear programming is NP-hard (Karp, 1972), and solvers will often solve the relaxed problem where the variables are allowed to be non-integer first. For the problem solved by Lpnet, this solution happens to be an integer solution most of the time. If there are non-integer entries, the running time will increase. For example, we observed this case for less than 1% of random distances for 80 taxa and Gurobi, and then the average CPU time was 307.27 sec. In an attempt to estimate the worst case, we constructed an example where the optimal solution of the relaxed problem contains no integer at all. For 80 taxa, Gurobi needed almost 14 hours for the solution. The distance matrix is available as no_integer.nex in the examples folder of our R package at https://github.com/yukimayuli-gmz/lpnet.

In summary, our Lpnet implementation can be used for up to 94 taxa, and with the Gurobi solver a solution will usually take at most a few minutes. The Rglpk solver works well for smaller instances but will struggle for more than 50 taxa.

We report running times for our heuristic in Table 8. Using a computer with more memory (128 GB), an example with 500 taxa took 28 hours.

**Table 8.** Heuristic method. User times for different number of taxa

| number of taxa | User time |
| --- | --- |
| 50 | 2.0631 sec |
| 100 | 42.5499 sec |
| 200 | 1176.228 sec |
| 280 | 5605.2504 sec |

All other CPU times reported in this article were obtained running Lpnet on a laptop with Windows 10 operating system, Intel Core i7-9750H 2.60 GHz CPU with 6 cores, and 16 GB of RAM.

## Data availability

The tree files, sequence alignments and distance matrices for the simulated data sets, as well as the distance matrices for the random distances are available at https://github.com/yukimayuli-gmz/data, while the real data set (DonoghueAll.nex) can be found in the examples folder of our R package.

